# Second-order boundaries segment more easily when they are density-defined rather than feature-defined

**DOI:** 10.1101/2023.07.10.548431

**Authors:** Christopher DiMattina

## Abstract

Previous studies have demonstrated that density is an important perceptual aspect of textural appearance to which the visual system is highly attuned. Furthermore, it is known that density cues not only influence texture segmentation, but can enable segmentation by themselves, in the absence of other cues. A popular computational model of texture segmentation known as the “Filter-Rectify-Filter” (FRF) model predicts that density should be a second-order cue enabling segmentation. For a compound texture boundary defined by superimposing two single-micropattern density boundaries, a version of the FRF model in which different micropattern-specific channels are analyzed separately by different second-stage filters makes the prediction that segmentation thresholds should be identical in two cases: (1) Compound boundaries with an equal number of micropatterns on each side but different relative proportions of each variety (*compound feature boundaries*) and (2) Compound boundaries with different numbers of micropatterns on each side, but with each side having an identical number of each variety (*compound density boundaries*). We directly tested this prediction by comparing segmentation thresholds for second-order compound feature and density boundaries, comprised of two superimposed single-micropattern density boundaries comprised of complementary micropattern pairs differing either in orientation or contrast polarity. In both cases, we observed lower segmentation thresholds for compound density boundaries than compound feature boundaries, with identical results when the compound density boundaries were equated for RMS contrast. In a second experiment, we considered how two varieties of micropatterns summate for compound boundary segmentation. In the case where two single micro-pattern density boundaries are superimposed to form a compound density boundary, we find that the two channels combine via probability summation. By contrast, when they are superimposed to form a compound feature boundary, segmentation performance is worse than for either channel alone. From these findings, we conclude that density segmentation may rely on neural mechanisms different from those which underlie feature segmentation, consistent with recent findings suggesting that density comprises a separate psychophysical ‘channel’.

## INTRODUCTION

Texture is an essential property of natural surfaces, and the visual perception of texture provides organisms with important information about surface material properties (**Motoyoshi, Nishida, Sharan, & Adelson, 2007; Komatsu & Goda, 2018**). Despite “texture” being hard to define precisely (**Bergen & Adelson, 1991; Adelson, 2001**), examination of various natural and man-made textures (e.g., **Brodatz, 1966; Olmos & Kingdom, 2004**) reveals that many textures are comprised of quasi-periodic spatial repetitions of smaller elements, often referred to as “micro-patterns” (**Julesz, 1981; Gurnsey, 1987; Zavitz & Baker, 2013; DiMattina & Baker, 2019**). One perceptually salient dimension along which textures can vary is the density of their constituent micropatterns. Indeed, in multidimensional scaling studies aimed at identifying the perceptual dimensions of texture, density was revealed to be one of the dimensions which explained a large proportion of variability in texture appearance (**Rao & Lohse, 1993, 1996**). Not only does density provide a useful cue for inferring material properties, but density gradients also provide powerful cues for inferring 3-D shape from 2-D images (**Cutting & Millard, 1984; Malik & Rosenholtz, 1997; Rosenholtz, 2014; Todd 2004**). Therefore, one should expect on purely ecological grounds that there should be a robust representation of texture density cues in biological visual systems. Indeed, studies using adaptation (**Sun, Kingdom, & Baker, 2017; Durgin & Proffitt, 1991, 1996; Durgin & Huk, 1997; Durgin, 2001**) and simultaneous density contrast (**Sun, Baker, & Kingdom, 2016**) have suggested that density is an independent dimension of texture to which the human visual system is highly sensitive. Furthermore, physiological work in nonhuman primates has also demonstrated neurons in V4 which exhibit tuning for texture density (**Hanazawa & Komatsu, 2001**), as well as neurons in other visual areas exhibiting sensitivity to texture density gradients (**Elmore, Rosenberg, DeAngelis, & Angelaki, 2019**).

In addition to texture providing information about the material composition of a single surface, textural differences between two different surfaces can be used for purposes of scene segmentation (**Landy & Graham, 2004; Landy, 2012; Victor et al., 2017; DiMattina & Baker, 2019**). Previous work has demonstrated that not only does density influence texture segmentation (**Zavitz & Baker, 2013**), but that density differences by themselves, in the absence of any other cues, can in fact enable segmentation (**Zavitz & Baker, 2014**). Assuming that texture micro-patterns are luminance-balanced, density segmentation cannot be explained by standard first-order visual mechanisms which simply compute regional luminance differences (e.g., Gabor filters). However, density segmentation can easily be explained by a two-stage model, commonly known as a Filter-Rectify-Filter (FRF) model (**Baker, 1999**; **Landy & Graham, 2004; DiMattina & Baker, 2019, 2021**), illustrated schematically in **Fig. 1**.

**Figure 1:**
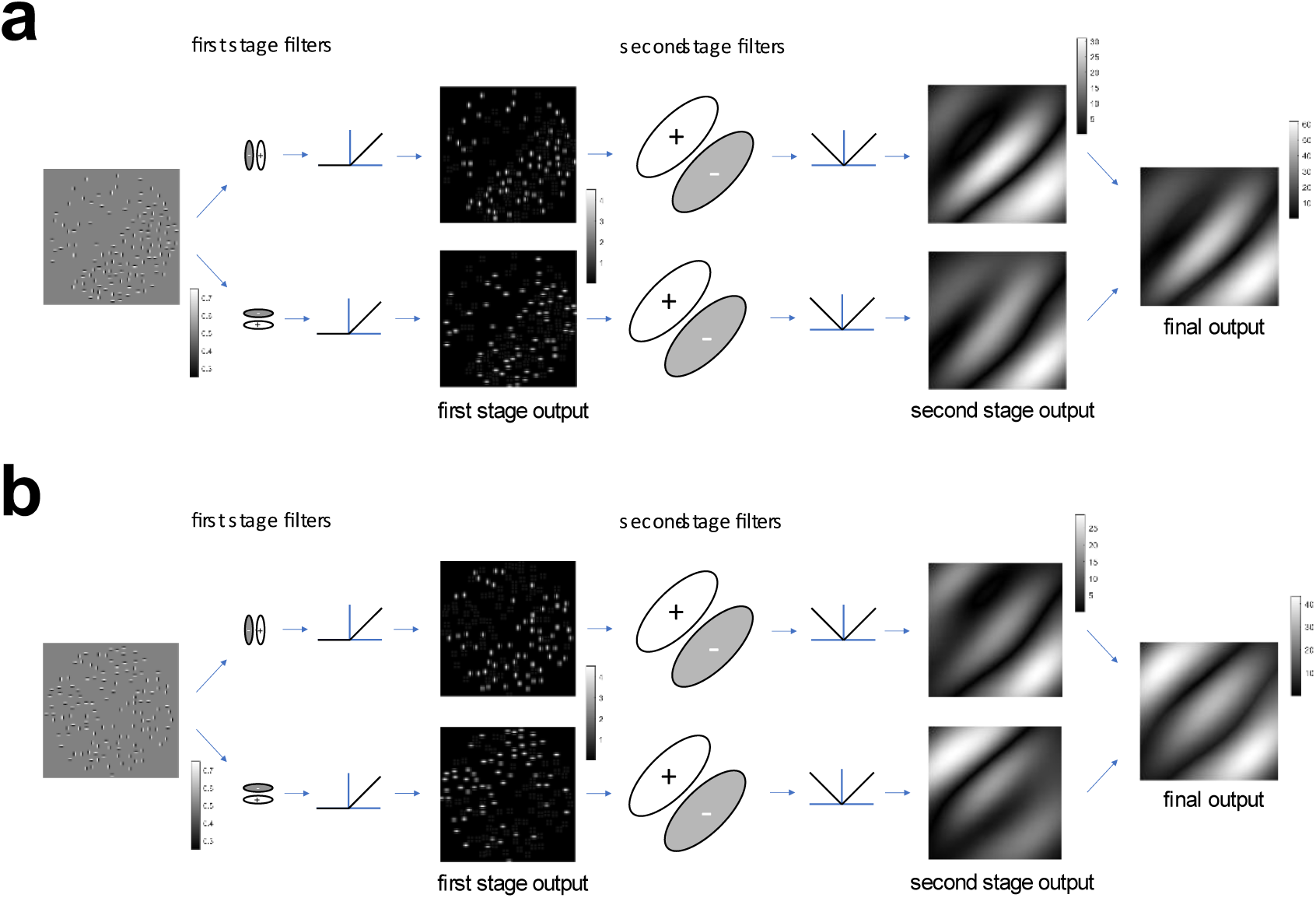
Illustration of Filter-Rectify Filter (FRF) model of second-order texture processing applied to compound density-defined and feature-defined boundaries. The original image (leftmost image) is first analyzed by Gabor filters defined at the spatial scale of the micro-patterns. The output of these first-stage filters is then half-wave rectified (here by the *relu()* function) to yield two images which represent the locations of each kind of micro-pattern (center-left images). These images are then analyzed on a global scale by a set of second-stage filters which look for differences in the density of each kind of micro-pattern on opposite sides of the boundary (center-right images). The full-wave rectified (here by the *abs()* function) outputs of these second-stage filters are then added to attain an output image (rightmost image). (a) FRF model applied to a compound density boundary comprised of two density boundaries (vertical and horizontal Gabors) super-imposed with the same phase of density modulation (0-degree phase shift). Note that there are peaks in the second-stage filter responses at the edges (where the image transitions from zero to non-zero contrast) as well as at the boundary (where the texture density changes). (b) FRF model applied to a compound feature boundary comprised of two density boundaries (vertical and horizontal Gabors) super-imposed with opposite phases of density modulation (180-degree phase shift). As in (a) there are peaks in the second-stage filter responses at the edges, as well as at the boundary where the texture pattern changes (feature boundary).

This FRF model (**Fig. 1**) contains a set of first-stage “local” filters defined on the spatial scale of the micro-patterns which detect the micro-patterns on each surface. The rectified output of these first-stage filters is then analyzed by much larger second-stage filters defined on the spatial scale of the texture boundary. If there are more micro-patterns of a particular kind on one side of the boundary than the other side, a second-stage filter will detect a difference in the outputs of the first-stage filters selective for that micro-pattern on opposite sides and will signal the presence of a boundary. In **Fig 1a**, we see a compound density boundary comprised of two different kinds of micro-patterns: horizontal and vertical Gabor functions. Each of these types of micro-patterns is detected by a different first-stage filter, giving rise to two separate channels of rectified first-stage output. These outputs are in turn analyzed by a set of channel-specific second-stage filters looking for either left-or right-oblique gradients in the first-stage filter outputs. As we see in **Fig. 1b**, this exact same FRF mechanism can also be used to segment compound feature boundaries with no density difference, but only differences in the relative proportion of two different kinds of micro-patterns on opposite sides of the boundary. This is because such boundaries are equivalent to two density boundaries (one for each micro-pattern type) superimposed in opposite-phase.

Although there are many possible variations on the basic FRF model architecture, in perhaps the most common form of the FRF model the outputs of different first-stage channels are analyzed by separate second-stage filters, as in the example shown in **Fig. 1** (**Malik & Perona, 1990; Bovik, Clark, & Geisler, 1990; Oruc & Landy 2002; Motoyoshi & Nishida, 2004; Motoyoshi & Kingdom, 2007; Zavitz & Baker, 2013; Victor 2017**). As we can see in **Fig. 1**, this version of the model would make the prediction that the compound feature and density boundaries should be about equally discriminable. However, at least subjectively, the compound density boundary (**Fig. 1a**) appears more obvious than the compound feature boundary (**Fig. 1b**), even though for both boundaries the proportion of each single micropattern on opposite sides is the same. This simple demonstration suggests the possible involvement of other mechanisms for compound density boundary segmentation that may not be utilized for compound feature boundaries.

In this paper, we compare feature-defined and density-defined second-order boundary segmentation for compound boundaries created by super-imposing two single-micropattern density boundaries. **Fig. 2a** illustrates compound boundaries defined by ON-center and OFF-center Difference-of-Gaussian (DOG) functions. When the two individual density boundaries are super-imposed with the same phase of density modulation, they create compound density boundary with equal numbers of each type of micro-pattern on each side, but different numbers of total patterns on opposite sides, as illustrated in **Fig. 2a** (top row). However, when the two single-micropattern density boundaries are super-imposed with opposite phases, they form a compound feature boundary, with the same number of total patterns on each side, but different numbers of each type on each side (**Fig. 2a**, bottom row). We compare the segmentation of compound feature and density boundaries as we systematically manipulate their visibility by varying the proportion of micropatterns of each type which are not balanced by one of the same kind on the opposite side of the boundary. These compound feature and density boundary stimuli are illustrated in **Fig 2b** for DOG micropatterns, and in **Fig. 2c** for horizonal and vertical Gabor micropatterns. We further demonstrate that the use of density as a segmentation cue cannot be attributed to changes in RMS contrast, as controlling for RMS contrast has no effect on segmentation thresholds for compound density boundaries.

**Figure 2:**
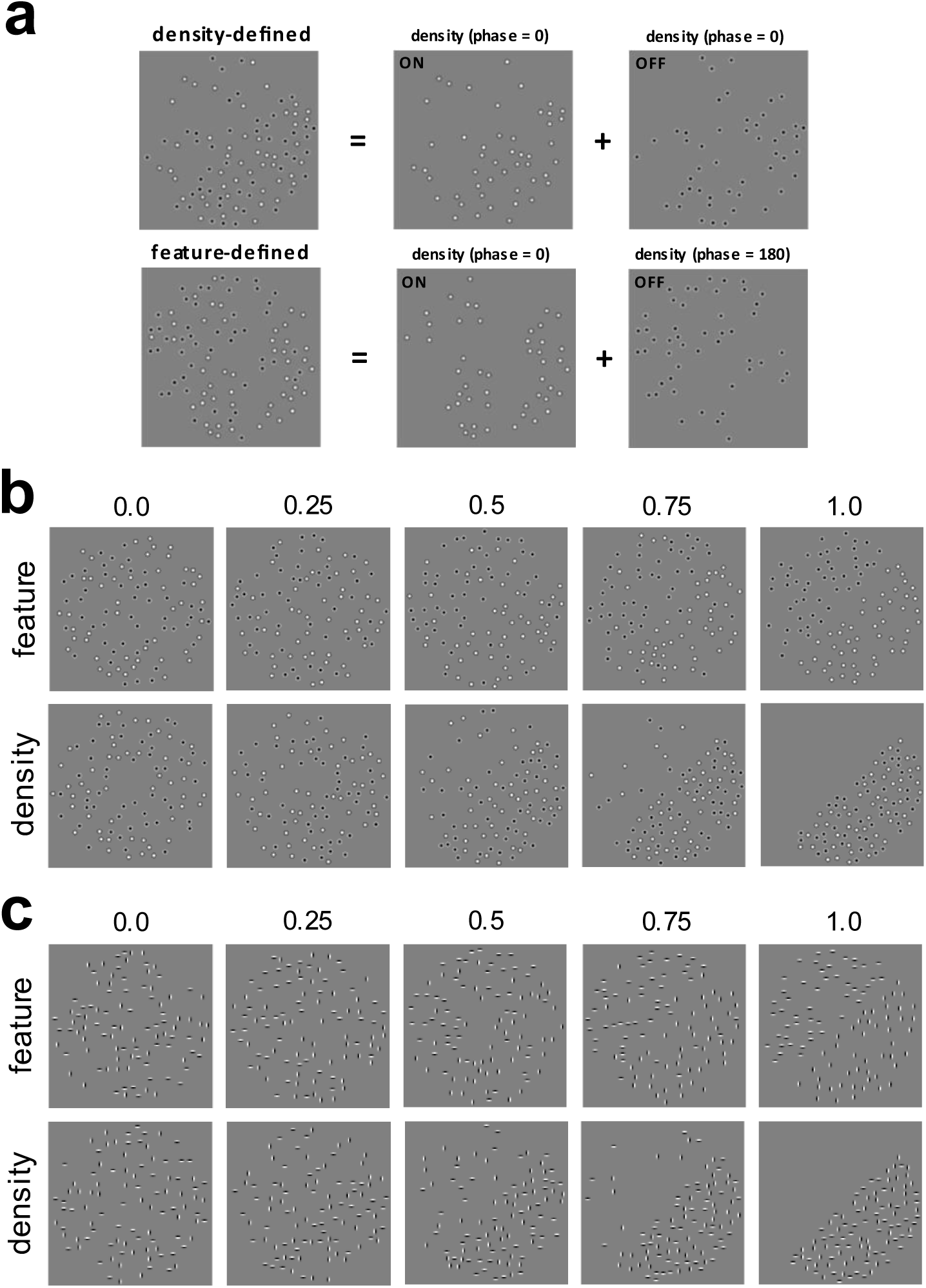
Illustration of compound feature-defined and density-defined boundaries, each of which are formed by combining single-micropattern density boundaries. (a) *Top:* A compound density-defined boundary is formed by combining two single-micropattern density boundaries having the same phase of density modulation (0-degree phase difference). Here the single-micropattern density boundaries are comprised of ON- and OFF-center DOG functions. *Bottom:* A compound feature-defined boundary is formed by combining two single-micropattern density boundaries having opposite density modulation phases (180-degree phase difference). (b) Demonstration of how compound boundary visibility is parameterized by varying the proportion 𝜋_𝑈_ of micro-patterns for each single-micropattern density boundary which are not balanced by a micropattern of the same kind on the opposite side. Here micro-patterns are ON- and OFF-center DOG functions. *Top*: Feature-defined boundary. *Bottom:* Density-defined boundary. (c) Same as (b), but for compound boundaries comprised of vertical (V) and horizontal (H) Gabor micropatterns.

In **Experiment 2**, we consider how single-micropattern density boundaries combine perceptually when they are superimposed to form compound density boundaries and feature boundaries. We find that when two single-micropattern density boundaries combine in-phase to form a compound density boundary (**Fig. 2a**, *top*), there is an improvement in psychophysical performance which is well described by a standard model of probability summation (**Kingdom, Baldwin, & Schmidtmann, 2015; Prins & Kingdom, 2018**). In other words, the whole is the (probability) sum of its parts. By contrast, when two single-micropattern density boundaries combine in opposite-phase to form a compound feature boundary (**Fig. 2a**, *bottom*), performance is worse for the compound feature boundary than for either of its constituent single-micropattern density boundaries. That is, the whole is less than the sum of its parts. This surprising and somewhat counter-intuitive result helps to provide strong constraints on possible computational models of texture segmentation and suggests that different mechanisms may be utilized for density and feature segmentation.

Despite several early studies investigating density as an important dimension of texture perception (**MacKay, 1964; MacKay, 1973; Anstis, 1974**), only somewhat recently has there been a re-emergence of interest in how texture density is represented by the visual system (**Durgin & Proffitt, 1991, 1996**). Recent discoveries using adaptation and simultaneous contrast have suggested that density may in fact be coded using multiple psychophysical “channels” (**Sun, Kingdom & Baker, 2016; Sun, Baker & Kingdom, 2017**), analogous to well-known psychophysical channels for orientation and spatial frequency (**DeValois & DeValois, 1988; Graham, 1989**). However, to the best of our knowledge, density segmentation has never been directly compared to feature segmentation within a single study. Furthermore, despite great interest in cue combination and summation models within psychophysics more generally (**Trommershauser, Kording, & Landy, 2011**), there has been little work systematically investigating how multiple types of micropattern combine for either density or feature boundary segmentation (**Saarela & Landy, 2012**; **DiMattina & Baker, 2019**). The current study rectifies these gaps in the literature, and furthermore provides powerful top-down constraints on computational theories of texture segmentation. We propose that additional insight into computational mechanisms may be obtained using psychophysical system identification methods recently developed in our laboratory (**DiMattina & Baker, 2019**), or behavioral decoding of deep neural networks trained on texture segmentation tasks (**Ronneberger et al., 2015**). Such modeling approaches could potentially provide more detailed insight into the underlying neural mechanisms of this perceptual phenomena.

## METHODS

### Stimuli and Task

#### Density and feature boundary stimuli

Compound density and feature boundaries comprised of two different kinds of micropatterns were formed by creating two different single-micropattern density boundaries having the same density modulation and then super-imposing them either in the same phase, or in opposite phase (**Fig. 2a**). As we can see in **Fig. 2a** (*top*), when the two single-micropattern density boundaries are super-imposed in-phase, the result is a *compound density boundary* having equal numbers of each kind of micro-pattern on each side of the boundary, but different numbers of total micro-patterns on opposite sides of the boundaries. Conversely, when the two single-micropattern density boundaries are super-imposed in opposite phase (**Fig. 2a**, *bottom*), this gives rise to a *compound feature boundary* having the same number of total micropatterns on each side of the boundary, but different proportions of each kind on opposite sides. Crucially, within each category of micro-pattern, the density difference is the same for both the compound feature and compound density boundaries. We refer to this class of texture stimuli comprised of two different kinds of micro-patterns as *compound micropattern textures* and have utilized them in previous work investigating luminance-based texture boundary segmentation (**DiMattina & Baker, 2021**; **DiMattina, 2022**). In this study, we considered the segmentation of two different types of compound micropattern textures, which formed either compound feature boundaries or compound density boundaries. The first type of compound micro-pattern texture we investigated was comprised of DC-balanced ON-center and OFF-center Difference-of-Gaussian (DOG) functions (**Fig. 2b**). Since all the individual micro-patterns were DC balanced, for both the compound feature and density boundaries there was no luminance difference on opposite sides of the boundary, so that all boundaries were second-order boundaries (**Baker, 1999**). The second type of compound micro-pattern texture we considered was comprised of horizontal and vertical odd-phase Gabor functions (**Fig. 2c**). Again, since each individual Gabor was DC balanced there was no net luminance cue across the boundary, making these boundaries second-order as well.

For both types of compound boundaries, we utilized a total of N = 48 micropatterns of each kind, with each micropattern defined in a 12-pixel square window. To speed up stimulus generation, micropatterns were pre-computed and on each trial placed in random non-overlapping locations within a circular window having a 320-pixel radius. For both compound density and feature boundaries, the absolute modulation phase was chosen at random on each trial to be either 0 (in-phase) or 180 (opposite-phase). This determined which side had more micro-patterns (in the case of compound density boundaries) or which side had more of each type of micro-pattern (in the case of compound feature boundaries).

The visibility of each of the two super-imposed boundaries was parameterized by the proportion 𝜋_𝑈_ of micro-patterns on each side that were not balanced by a micropattern of the same kind on the opposite side of the boundary. With this parameterization, 𝜋_𝑈_ = 0 corresponds to every micro-pattern being balanced (i.e., same number on each side), and 𝜋_𝑈_ = 1 corresponds to every micro-pattern being unbalanced (i.e., all the micropatterns on one side, and none on the other side). This parameterization is illustrated in **Fig. 2b, c** for the two different types of compound micropattern textures used in this study.

#### Psychophysical task

Observers performed a single-interval classification task (**Fig. 3a**) identifying the orientation of a boundary (feature-defined or density-defined) as either right-oblique (+45 deg. with respect to vertical) or left-oblique (−45 deg. with respect to vertical). This *orientation-identification* task has been used in several previous studies investigating boundary segmentation (**Arsenault, Yoonessi, & Baker, 2011**; **Zavitz & Baker, 2013, 2014; DiMattina & Baker, 2019, 2021**). Observers were given feedback after each trial to maximize task performance.

**Figure 3:**
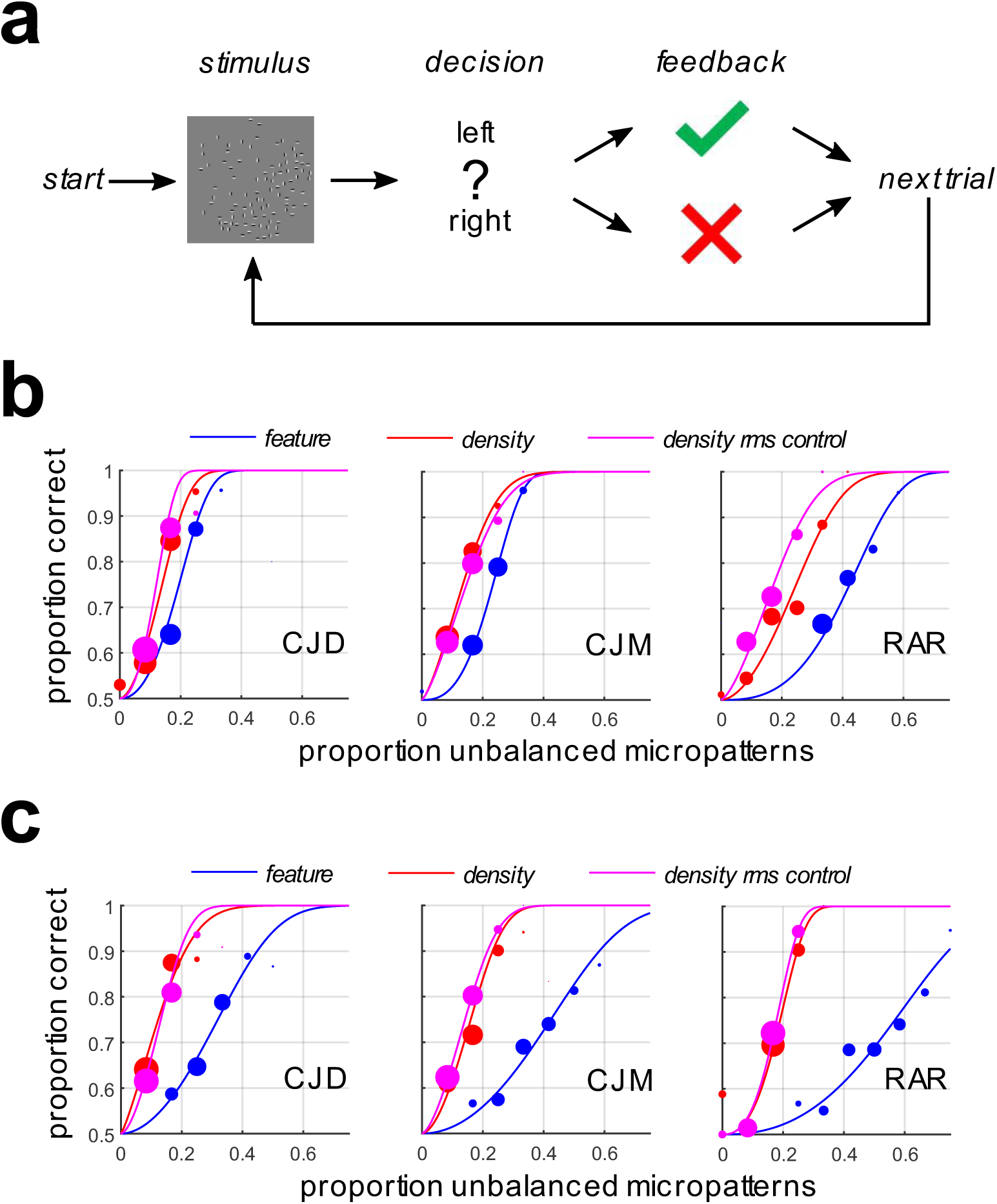
Segmentation thresholds for density-defined and feature-defined boundaries. (a) Illustration of the orientation identification task. Compound feature-defined and density-defined boundaries having varying degrees of visibility were presented with a left-oblique or right-oblique orientation. (b) Psychometric functions for observers segmenting compound feature-defined boundaries (blue symbols + curves), compound density-defined boundaries (red symbols + curves), and RMS-equated compound density-defined boundaries (magenta symbols + curves). Here the constituent single-micropattern density boundaries were comprised of ON- and OFF-center DOG functions (**Fig 2b**). (c) Same as (b) but for compound feature- and density-defined compound boundaries comprised of horizontal (H) and vertical (V) Gabor micropatterns (**Fig. 2c**).

#### Observers

Author CJD and two naïve undergraduate student researchers (CJM, RAR) served as observers in these experiments. All observers had normal or corrected-to-normal visual acuity. All observers gave informed consent, and all experimental procedures were approved by the FGCU IRB (Protocol 2014-01), in accordance with the Declaration of Helsinki.

#### Visual Displays

Stimuli were presented in a dark room on a 1920×1080, 120 Hz gamma-corrected Display++ LCD Monitor (Cambridge Research Systems LTD®) with mid-point luminance of 100 cd/m^2^. This monitor was driven by an NVIDA GeForce® GTX-645 graphics card, and experiments were controlled by a Dell Optiplex® 9020 running custom-authored software written in MATLAB® making use of Psychtoolbox-3 routines (**Brainard, 1997; Pelli, 1997**). Observers were situated 133 cm from the monitor using a HeadSpot® chin-rest, so that the 320×320 pixel stimuli subtended approximately 5 deg. of visual angle.

### Experimental Protocol

#### Experiment 1

The objective of Experiment 1 was to simply compare the segmentation thresholds for compound feature- and density-defined boundaries in units of the proportion of unbalanced micropatterns of each of their constituent single-micropattern boundaries. Boundary visibility (parametrized by the proportion 𝜋_𝑈_ of unbalanced micropatterns) was adjusted for each type of stimulus using a 1-up, 2-down staircase procedure, which focuses trials at stimulus levels leading to 70.71% correct performance (**Leek, 2001**). The staircase was defined on a set of stimulus levels varying from 𝜋_𝑈_ = 0 to 𝜋_𝑈_ = 1 in N = 13 evenly spaced steps. Observers first completed a practice set of trials which was not analyzed, and then did two more sets of N = 240 trials each, which were then combined for analysis.

Since our stimuli were DC-balanced, modulation of density did not give rise to a luminance difference. However, it is still the case that modulating the density will give rise to differences in RMS contrast, and it is well known that contrast is a second-order cue which supports segmentation (**Dakin & Mareschal, 2000**). Therefore, as a control, we ran an additional condition for the density-defined stimuli in which we equalized the RMS contrast on both sides of the boundary by re-scaling the micropattern amplitudes as we varied density.

#### Experiment 2

A large body of work in perception science has investigated how multiple stimuli combine to reach threshold, with a strong emphasis on determining mathematical rules for cue combination (**Trommershauser, Kording, & Landy, 2011**). Our compound micro-pattern textures lend themselves to cue combination studies naturally since they are comprised of two non-overlapping single-micropattern density boundaries. To determine whether the cue-combination rule depends on whether the single micro-pattern boundaries are aligned in-phase to form a compound density boundary (**Fig. 2a**, top) or opposite-phase to form a compound feature boundary (**Fig. 2a**, bottom), we performed a cue-combination experiment in which we obtained psychometric functions for each cue by itself and both cues together. For this experiment, we varied the proportion of unbalanced micropatterns and presented either single-micropattern density boundary (A), the complementary single-micropattern density boundary (B) or both boundaries together (A + B), to form either a compound density boundary (**Fig. 2a**, top) or compound feature boundary (**Fig. 2a**, bottom). Levels were 7 evenly spaced steps from 𝜋_𝑈_ = 0 to 𝜋_𝑈_ = 0.5, and stimuli were presented using the method of constant stimuli with N = 80 trials at each level for each of the three conditions (A, B, A + B). Data was collected over a series of 4 different blocks of trials, each of which contained N = 20 trials at each level for each of the three conditions.

### Data Analysis

#### Psychometric function fitting and bootstrapping

Although this experiment is a single-interval task, one can model the psychophysical data using the same formalism developed for a two-interval task if one assumes that a decision variable is formed by subtracting the outputs of a mechanism looking for a left-oblique boundary from one looking for a right-oblique boundary. To model our data, we made use of a signal-detection-theory (SDT) psychometric function modeling framework utilized in several previous studies (**Kingdom et al., 2015; Prins & Kingdom, 2018**). In the STD framework, the proportion of correct responses for a stimulus having level 𝑥 is given by the expression

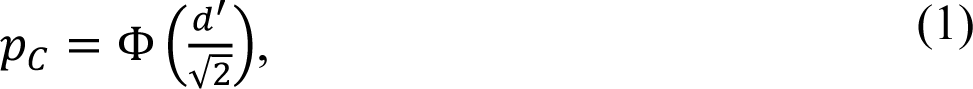

where the stimulus discriminability 𝑑^′^ = [𝑔𝑥]^𝛼^ is given by a power-law transducer function of the stimulus level with gain 𝑔 and exponent 𝛼. This function was fit to psychophysical trial data by numerically optimizing the transducer function parameters to maximize the data likelihood, making use of MATLAB routines available at www.palamedestoolbox.org. From these fits, we could then obtain an estimate for the just-noticeable difference (JND) threshold, which we took as the stimulus level 𝑥_𝐽_at which performance was 75% correct. This threshold could be obtained from Eq. (1) by setting 𝑝_𝐶_ = 0.75 and making use of the fact that the normal CDF Φ is invertible to solve for

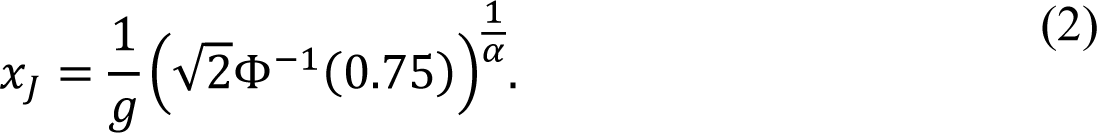

To obtain confidence intervals for our threshold estimates, we made use of a bootstrapping procedure (**Efron & Tibshirani, 1994**) to obtain a distribution of thresholds using N = 1000 resampled datasets to which we fit our psychometric function (1).

#### Summation model

In order to better understand how observers process compound density boundaries comprised of two super-imposed single-micropattern density boundaries, we fit two standard summation models to one of the conditions in Experiment 2 (**Kingdom et al., 2015**). The first model, known as probability summation (PS), assumes that there exist distinct mechanisms for processing each boundary cue, and that detection of the compound stimulus occurs when either mechanism (or both) reaches its threshold. The second model, known as additive summation (AS), assumes that there exists a common mechanism which additively combines inputs from distinct mechanisms sensitive to each boundary cue, and that detection of the compound stimulus occurs when this common mechanism reaches threshold. Therefore, with AS (but not with PS), it is possible for sub-threshold summation to occur. That is, even if each individual cue is below threshold for its dedicated mechanism, the combined effects of both cues presented together may exceed threshold for the common mechanism, enabling detection. Here we compare probability summation to a simple linear additive summation model used in previous work (**Kingdom et al., 2015**). MATLAB routines for fitting both models are available online at www.palamedestoolbox.org.

## RESULTS

### Experiment 1

Given two single-micropattern density boundaries, one may combine then either in-phase (**Fig. 2a**, top) or opposite-phase (**Fig. 2b**, bottom) to form compound density boundaries or compound feature boundaries. Compound density boundaries have a different number of micropatterns on opposite sides, but the same number of both kinds of micropattern on each side. Conversely, compound feature boundaries have the same number of total micropatterns on opposite sides, but different proportions of the two kinds of micropattern on each side. For both the compound density and feature boundaries, we vary their visibility by varying for each of their constituent single-micropattern boundaries the proportion 𝜋_𝑈_ of micro-patterns on each side that are not balanced by a micro-pattern of the same kind on the opposite side (**Fig. 2b**, **c**). This is illustrated in **Fig. 2b** for compound boundaries (top: feature, bottom: density) formed by combining single-micropattern density boundaries comprised of DC-balanced ON-center difference of gaussian (DOG) functions, and DC-balanced OFF-center DOG functions. Likewise, this is illustrated in **Fig. 2c** for composite boundaries (top: feature, bottom: density) comprised of single-micropattern density boundaries comprised of DC-balanced vertical Gabor functions and horizontal Gabor functions.

We observed that when we vary the visibility of each single-micropattern density boundary by varying 𝜋_𝑈_, the visibility of compound density-defined boundaries was much stronger than that of compound feature-defined boundaries, as measured by segmentation thresholds in a boundary orientation-identification task (**Fig. 3a**). **Fig 3b** shows the psychometric functions obtained from three observers segmenting compound feature-defined (blue curves) and density-defined boundaries (red curves) comprised of complementary ON/OFF DOG functions (**Fig. 2b**). We find significantly lower segmentation thresholds for the density-defined boundaries, with 95% confidence intervals of the difference (feature-density) given in **Table 1**. Critically, these confidence intervals (based on N = 1000 bootstraps) do not contain zero.

**Table 1:** Difference between segmentation thresholds for feature-defined and density-defined composite boundaries comprised of ON- and OFF-center DOG functions (**Fig. 2b**). Median differences and 95% confidence intervals of the difference were computed from N = 1000 bootstraps.

| Observer | Median diff. | 95% CI |
| --- | --- | --- |
| CJD | 0.0581 | [0.0195, 0.1817] |
| CJM | 0.0981 | [0.0631, 0.1291] |
| RAR | 0.1742 | [0.1206, 0.2292] |

Qualitatively similar results were obtained in the case of compound density-defined and feature-defined boundaries comprised of horizontal and vertical Gabor micropatterns (**Fig. 2c**), as illustrated in **Fig. 3c** (blue curves: feature-defined, red-curves: density-defined). **Table 2** summarizes the median difference between thresholds for the two kinds of boundaries and 95% bootstrapped confidence intervals of the difference.

**Table 2:**
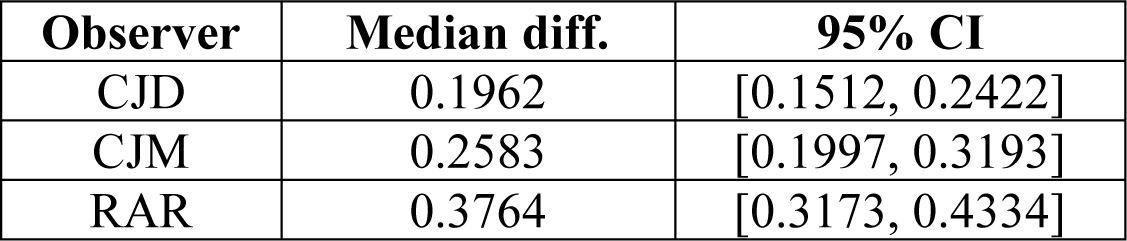
Same as **Table 1** except for feature-defined and density-defined composite boundaries comprised of horizontal and vertical Gabor functions (**Fig. 2c**).

For both classes of stimuli (DOG/Gabor), all the micropatterns were DC balanced so that varying density would not give rise to a luminance difference across the boundary which could be used for segmentation. However, even with this precaution, one unavoidable consequence of varying micropattern density is that one also varies the RMS contrast, which is defined for a texture as the standard deviation of the pixel intensities divided by the mean pixel intensity (**Zavitz & Baker, 2013**). To ensure that the reduced segmentation thresholds for compound density boundaries relative to compound feature boundaries were not a trivial consequence of the confound between density and contrast, we re-scaled the Michelson contrast of the micropatterns on each side of the boundary to equate the RMS contrast as we varied density. This resulted in nearly identical results as the compound density boundaries which were not RMS equated, as we see in **Fig. 3b** and **Fig. 3c** (magenta lines). As we see in **Tables 3** and **4**, our results remain unchanged for these RMS-equated compound density boundaries (compare to **Tables 1** and **2**, respectively).

**Table 3:** Same as **Table 1** but for RMS-equated compound density boundaries.

| Observer | Median diff. | 95% CI |
| --- | --- | --- |
| CJD | 0.0736 | [0.0353, 0.1111] |
| CJM | 0.0876 | [0.0511, 0.1216] |
| RAR | 0.2482 | [0.1992, 0.2975] |

**Table 4:** Same as **Table 2** but for RMS-equated compound density boundaries.

| Observer | Median diff. | 95% CI |
| --- | --- | --- |
| CJD | 0.1827 | [0.1349, 0.2270] |
| CJM | 0.2818 | [0.2322, 0.3396] |
| RAR | 0.3889 | [0.3258, 0.4467] |

In fact, directly comparing segmentation thresholds for the standard density boundaries and RMS equated density boundaries only revealed a significant difference in the case of a single observer (RAR), and only for the DOG micropattern stimulus, as summarized in **Table 5** and **Table 6**. Thresholds for all observers and conditions are summarize in **Fig. 4**.

**Figure 4:**
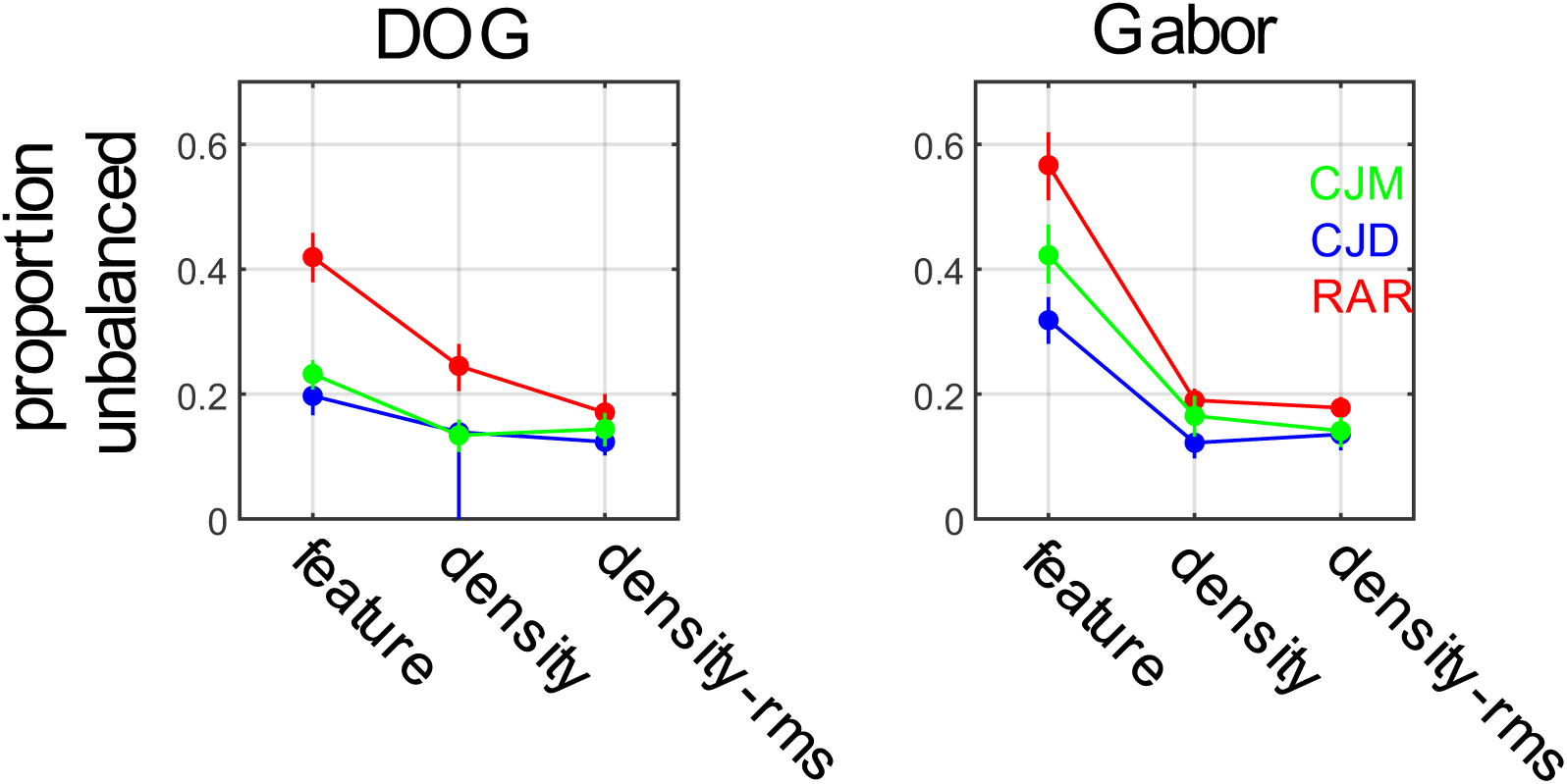
Segmentation thresholds for feature-defined, density-defined, and RMS-equated density defined compound boundaries for stimuli comprised of ON/OFF DOG functions (*left panels*) and H/V Gabor functions (*right panels*).

**Table 5:** Median difference between segmentation threshold for compound density boundaries comprised of ON/OFF DOG functions with and without equated RMS contrast.

| Observer | Median diff. | 95% CI |
| --- | --- | --- |
| CJD | 0.0165 | [-0.1031, 0.0458] |
| CJM | -0.0105 | [-0.0480, 0.0250] |
| RAR | 0.0746 | [0.0205, 0.1234] |

**Table 6:** Same as **Table 5** but for compound density boundaries comprised of H/V Gabors.

| Observer | Median diff. | 95% CI |
| --- | --- | --- |
| CJD | -0.0140 | [-0.0493, 0.0220] |
| CJM | 0.0240 | [-0.0178, 0.0636] |
| RAR | 0.0115 | [-0.0170, 0.0385] |

### Experiment 2

Given that the compound density and feature boundaries are each comprised of two non-overlapping single-micropattern density boundaries, it is of great interest to understand how these two single-micropattern density boundaries combine, and whether the rules by which they combine differ when they combine to form compound density or compound feature boundaries. To investigate this systematically, we compared segmentation thresholds for density boundaries comprised of one micropattern type (A), the other micropattern type (B), and both micropatterns together (A + B), which were combined in-phase to form a compound density boundary or in opposite phase to form a compound feature boundary.

As we see in **Fig. 5** (leftmost column), for compound density boundaries constituted from ON- and OFF-center DOG functions, we see that for all three observers performance for the compound boundary (black symbols) is better than that of either of the individual boundaries (blue symbols, green symbols). Error-bars demonstrate the 95% binomial proportion confidence intervals for the compound stimulus. The center-left column shows the difference between the proportion of correct responses for the composite stimulus and each of the individual density boundaries, with error bars showing the 95% confidence interval of the difference (binomial proportion). As we see, performance for the compound stimulus is significantly better than that of either of the constituent single-micropattern density boundaries. Fitting two popular models of sensory cue combination (probability summation, additive summation: see METHODS) to the data reveals a better fit to the data (larger log-likelihood) by the probability summation model (**Table 7**). That is, when two density boundaries combine to form a compound density boundary, the whole is the (probability) sum of its parts.

**Figure 5:**
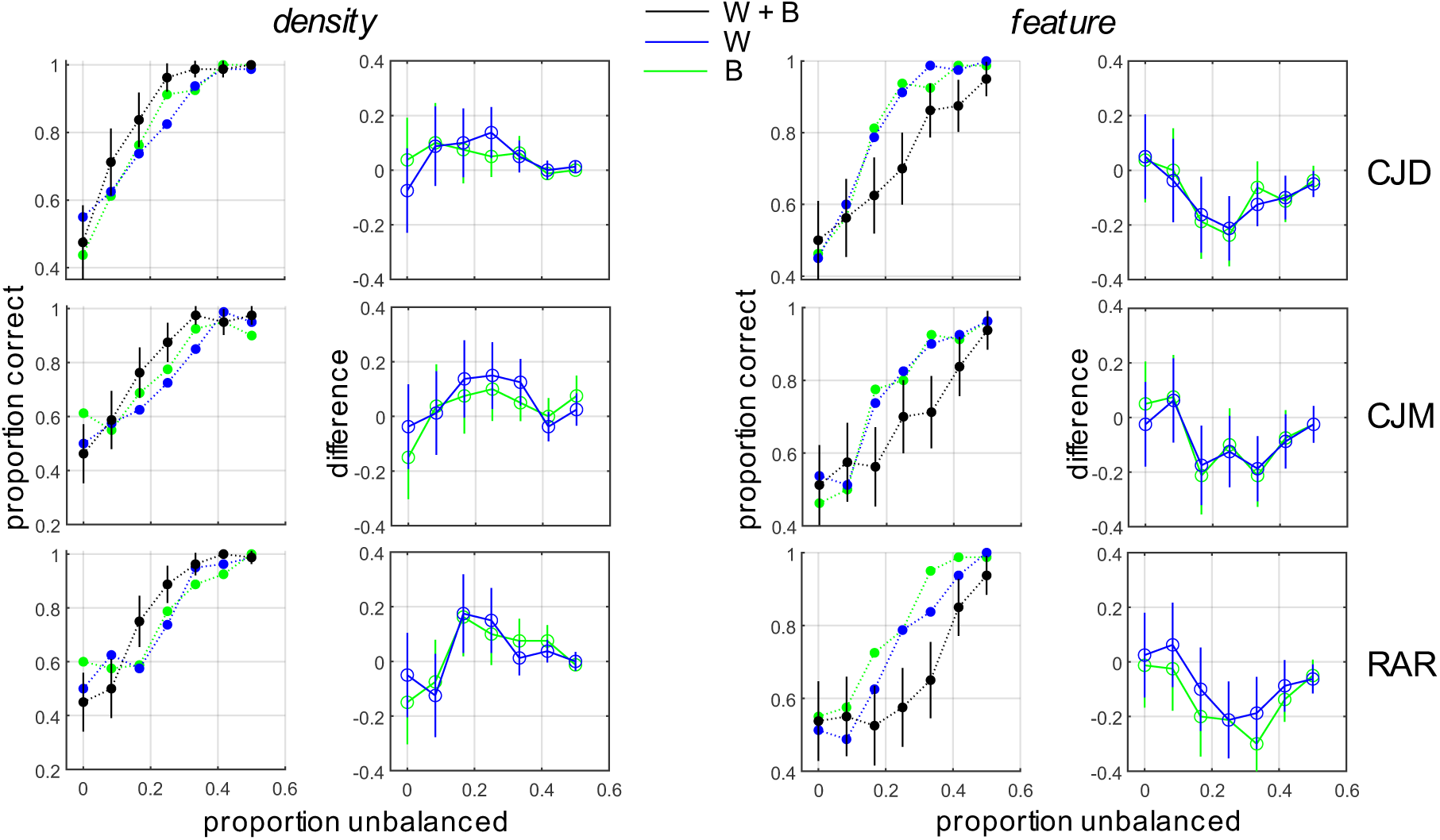
Segmentation performance for density-defined (left two columns) and feature-defined compound boundaries (right two columns). *Left:* Psychometric functions for single-micropattern boundaries (blue and green symbols) and the density-defined compound boundary (black symbols). Black lines indicate 95% binomial proportion confidence intervals. *Center-left:* Difference in segmentation performance between the composite density-defined boundary and each of the single-micropattern boundaries. Lines indicate 95% binomial proportion confidence intervals of the difference. *Center-right:* Psychometric functions for single-micropattern boundaries (blue and green symbols) and the composite feature-defined boundary (black symbols). *Right:* Difference in segmentation performance between the composite feature-defined boundary and each of the single-micropattern boundaries.

**Table 7:** Log-likelihood of probability summation models and additive summation models fit to density segmentation experiment for compound density boundaries comprised of ON/OFF center DOG functions.

| Observer | PS | AS |
| --- | --- | --- |
| CJD | -585.458 | -592.781 |
| CJM | -761.489 | -771.063 |
| RAR | -710.442 | -716.705 |

By contrast, we see from the center-right column of **Fig. 5** that when two single-micropattern boundaries combine out of phase to form a compound feature boundary, that performance for the compound feature boundary (black symbols) is worse than that for the individual single-micropattern density boundaries that comprise it (blue, green symbols). Furthermore, the difference in performance is significant, as evidenced by the 95% confidence intervals of the difference (binomial proportion) not containing zero. Hence, instead of summing, the two cues interfere with each other, so that performance for the whole is significantly worse than that for either of its parts (**Fig. 5**, rightmost column).

Similar results were obtained for obtained for compound density-defined boundaries and compound feature boundaries comprised of horizontal and vertical Gabor micro-pattern density boundaries (**Fig. 6**, **Table 8**). As with the DOG micropattern stimuli, when the H and V density boundaries combine in-phase to form a new compound density boundary, performance improves significantly, and the combination rule is best explained by probability summation (**Kingdom et al., 2015**). Conversely, when they combine in opposite phase to form a compound feature boundary, performance for the whole is much worse than either of its individual parts (**Fig. 6**, rightmost column).

**Figure 6:**
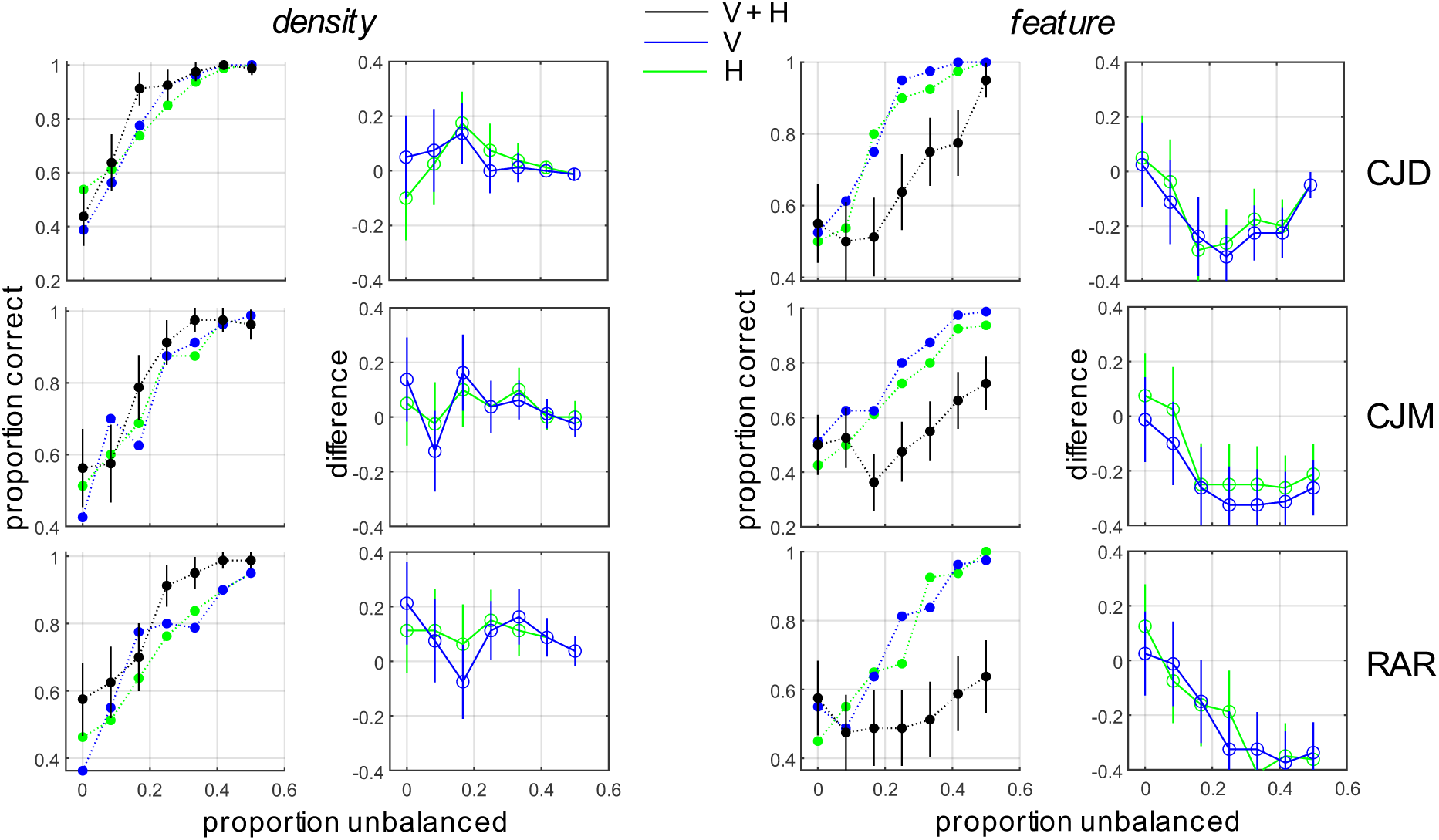
Same as **Fig. 5** except for density-defined and feature-defined boundaries comprised of H/V Gabor micropatterns (**Fig. 2c**).

**Table 8:** Same as **Table 7**, but for compound density boundaries comprised of H/V Gabor functions.

| Observer | PS | AS |
| --- | --- | --- |
| CJD | -576.094 | -590.959 |
| CJM | -690.971 | -707.733 |
| RAR | -774.710 | -776.246 |

## DISCUSSION

### Density as a perceptually important texture dimension

Although it is difficult to completely specify a set of intuitive, verbally describable features which fully capture the perceptual dimensions of visual texture (**Rao & Lohse, 1996; Bhushan et al., 1997; Cho, Yang & Hallett 2000**), any such specification will inevitably include density or closely related concepts (for instance, fineness/coarseness or granularity). This is because natural visual textures typically involve quasi-periodic spatial repetition of local patterns defined on a smaller spatial scale than the entire surface. In a series of seminal exploratory studies on the perceptual dimensions of texture, **Rao & Lohse** (**1993**, **1996**) had human observers rate the similarity of natural and man-made textures from the Brodatz album (**Brodatz, 1966**) and then performed statistical analyses on the resulting similarity ratings to determine which verbally describable texture dimensions best describe the variability observed in natural texture appearance. They found that in their set of 12 verbal descriptors (which included “density”), that only 1 descriptor explained less than 5% of the variability and no descriptor explained more than 10%, with density explaining 8.29% (**Rao & Lohse, 1996)**. Principal components analysis revealed four significant principal components, with density loading significantly onto the third principal component (**Rao & Lohse, 1996**).

Additional evidence for the perceptual importance of texture density for texture appearance comes from work by **Kingdom, Hayes & Field (2001)** in which they compared the visual discriminability of Gabor micropattern textures differing in various higher-order moments of the distributions from which their contrasts were drawn. They found that human observers were most sensitive to differences in the 4^th^ moment or kurtosis, which determines the texture density/sparsity, and less sensitive to differences in histogram variance (2^nd^ moment) or skew (3^rd^ moment).

In addition to its importance for texture appearance, density can influence texture segmentation, and even serve as a segmentation cue in the absence of other information. Early work demonstrated that density improves segmentation performance, with structures with discriminable elements segmenting more easily when these elements are closely spaced (**Nothdurft, 1985**). More recently, phase-scrambling natural textures was shown to improve performance on a contrast-based segmentation task using natural textures since phase-scrambling increases the texture density (**Arsenault, Yoonessi, & Baker, 2011**). Increasing micropattern density was also shown to lower segmentation thresholds when textures were segmented using micropattern orientation or spatial frequency cues (**Zavitz & Baker, 2013**). Finally, a third study from this line of work demonstrated that density in and of itself was adequate as a segmentation cue (**Zavitz & Baker, 2014**), a result which we replicate here (**Fig. 3**).

### Mechanistic bases of texture density representation

Adaptation is often referred to as the psychologist’s microelectrode because it can reveal the existence of neuronal populations tuned to sensory dimensions of interest (**Frisby, 1980**). Although early adaptation studies into the representation of density suffered from problems of confounds between spatial frequency content and density (**Antsis, 1974**), better-controlled studies revealed that density adaptation effects could not be explained as a simple consequence of spatial frequency adaptation (**Durgin & Proffitt, 1991, 1996**; **Durgin & Huk, 1997**). Evidence that density is represented independently of simple dimensions like orientation and spatial frequency comes from an adaptation study in which it was shown that density after-effects transfer when the adapting and test micropatterns are different, although the effect is stronger when the types of micropatterns are matched (**Durgin & Huk, 1997**). Unlike the case with luminance and contrast adaptation, density adaptation and simultaneous density contrast are binocular in nature (**Durgin, 2001**), suggestive of at least some mechanisms for density coding which are invariant to these low-level correlates of density. Our current results corroborate these findings, as we see that density-based segmentation is unaffected by controlling for RMS contrast (**Fig. 3, 4**). While density-based segmentation cannot be explained as a simple consequence of changes in contrast or luminance, it is the case that there are some mechanisms which are simultaneously sensitive to contrast and density, as revealed by recent summation studies (**Morgan, MacLeod, & Solomon, 2022**).

Although earlier work on density adaptation and simultaneous density contrast suggested that these effects were uni-directional, more recent work has demonstrated both phenomena are bi-directional (**Sun, Baker, & Kingdom, 2016**; **Sun, Kingdom, & Baker, 2017**). This bi-directionality of these contextual effects is consistent with the notion of density “channels” tuned to various ranges of densities. If there are psychophysical channels sensitive to density, it might suggest the existence of populations of neurons tuned to different ranges of densities. To our knowledge, this has not been directly investigated in neurophysiology studies, although several studies have demonstrated a general selectivity for textures in higher-level ventral stream areas (**Hanazawa & Komatsu, 2001**; **Okazawa, Tajima, & Komatsu, 2015, 2017**). Perhaps the closest study to directly address the issue of whether neurons were tuned for texture density in the frontal-parallel plane was the work by (**Hanazawa & Komatsu, 2001**) which demonstrated neurons in V4 were sensitive to the spacing between micro-patterns (i.e., density) in artificial texture stimuli. However, a systematic investigation of tuning for density was not the main focus of this work, and therefore the nature of the population code for density remains uncertain. Another body of neurophysiology literature focused on 3-D vision has investigated neural tuning to density gradients, finding neurons in areas V3A and CIP sensitive to various combinations of slant and tilt giving rise to density gradients of a checkered stimulus (**Elmore, Rosenberg, DeAngelis, & Angelaki, 2019**). While these studies are focused on a different set of questions than the psychophysical literature on density encoding, they are strongly suggestive of the existence of neuronal populations in multiple visual areas which may also encode density in the frontal-parallel plane.

### Novel contributions and future directions

The binary micropattern textures used here (**Fig. 2**) and in previous work (**DiMattina & Baker, 2021**; **DiMattina, 2022**) present an opportunity to ask several interesting and novel questions about the representation of density in biological visual systems. With our stimuli, one can directly compare the segmentation of density and feature boundaries in which the “signal” from each variety of micropattern is identical in both cases, but their relative spatial phase is varied (**Fig. 2**). Doing this, we find that a standard FRF model of second-order boundary detection (**Fig. 1**) is not adequate to simultaneously explain the segmentation of density-defined and feature-defined boundaries, as it would predict identical segmentation thresholds for both, contrary to our findings (**Fig. 3**, **4**). Furthermore, in the case where density boundaries are comprised of multiple species of micro-patterns, we can use our binary micropattern stimuli to determine the rules by which summation takes place across micro-pattern types, which can in turn inform computational models of texture segmentation (see **DiMattina & Baker, 2019**). Neither of these interesting questions about density representation can be addressed with single species micropatterns like those used in nearly all previous studies of density representation (**Durgin & Proffitt, 1991, 1996**; **Durgin & Huk, 1997**; **Sun, Baker, & Kingdom, 2016**; **Sun, Kingdom, & Baker, 2017**; **Morgan, MacLeod, & Solomon, 2022**). Therefore, we feel that the binary micropattern textures presented here provide a novel and useful tool for studies of density representation.

Despite a large body of work on density representation in general, relatively little work has directly considered density as a segmentation cue (**Zavitz & Baker, 2014**). To our knowledge, this is the first study to directly compare density segmentation with feature segmentation in a controlled manner, showing quite compellingly that density-defined boundaries segment more easily. Although the goal of the present paper is to present qualitative description of a novel finding which cannot be easily accounted for by one common form of the FRF model (**Malik & Perona, 1990; Bovik, Clark, & Geisler, 1990; Oruc & Landy 2002; Motoyoshi & Nishida, 2004; Motoyoshi & Kingdom, 2007; Zavitz & Baker, 2013; Victor 2017**), there are several plausible FRF style-models of segmentation which can potentially explain our results. One simple interpretation of our results is that each species of micro-pattern is analyzed by two populations of neurons first-stage filters: (1) Filters that respond to one micro-pattern only and (2) Filters that respond equally well to both micro-patterns. In the case of density boundaries, both populations provide “signal” which can be utilized by a second-stage filter for the segmentation task. However, in the case of the feature-defined boundaries, Population 1 provides “signal” to a second-stage filter, whereas Population 2 provides “noise”. Therefore, according to this conceptual model we should expect lower segmentation thresholds for density-defined boundaries than feature-defined boundaries. Quantitative implementation of this conceptual model using psychophysical system identification experiments (**DiMattina & Baker, 2019**) will be an interesting avenue for future work.

Although the FRF model has been widely used as a phenomenological model of performance in texture-segmentation tasks, it lacks biological plausibility in the sense that it goes from being completely “blind” to texture boundaries in the first stage to detecting them in a single stage (second stage filters) which “see” the whole stimulus. By contrast, available neurophysiology evidence suggests that there is a gradual emergence of texture selectivity over a number of processing stages (**Okazawa, Tajima, & Komatsu, 2015, 2017**), which is sensible given the gradual increase in receptive field sizes. Therefore, a better computational modeling scheme for relating perception to neural mechanisms may be decoding deep neural networks (**Kriegeskorte 2015**; **Serre, 2019**) trained on object recognition tasks (**Krizhevsky, Sutskever, Hinton, 2017**), or trained directly on texture segmentation tasks (**Ronneberger et al. 2015**). Early work in this field demonstrated that even when networks are trained on a generic task like object recognition, units in intermediate layers often learn selectivity for textures (**Zeiler & Fergus, 2014**). Developing models which decode these texture-sensitive layers may be a fruitful approach to modeling human performance in texture segmentation tasks like those considered here. It is of great interest for future research to see if biologically plausible deep neural network models exhibit the same preference for compound density boundaries over compound feature boundaries which we have demonstrated here.

## Acknowledgements

We thank FGCU undergraduates in the Computational Perception Lab for help with data collection.

